# Aging-related iron deposit prevents the benefits of HRT from late postmenopausal atherosclerosis

**DOI:** 10.1101/2022.06.24.497502

**Authors:** Tianze Xu, Jing Cai, Lei Wang, Li Xu, Hongting Zhao, Fudi Wang, Esther Meyron-Holtz, Fanis Missirlis, Tong Qiao, Kuanyu Li

## Abstract

Postmenopausal atherosclerosis has been attributed to estrogen deficiency. The beneficial effect of hormone replacement therapy (HRT), however, is lost in late postmenopausal women with atherogenesis. We asked whether aging-related iron accumulation affects estrogen receptor α (ERα) expression explaining HRT inefficacy. A negative correlation between aging-related systemic iron deposition and ERα expression in postmenopausal AS patients was established. In an ovariectomized ApoE^-/-^ mouse model, estradiol treatment had contrasting effects on ERα expression in early versus late postmenopausal mice. ERα expression was inhibited by iron treatment in cell culture and iron-overloaded mice. Combined treatment with estradiol and iron further decreased ERα expression, mediated by iron-regulated E3 ligase Mdm2. In line with these observations, cellular cholesterol efflux was reduced and endothelial homeostasis was disrupted and, consequently, atherosclerosis was aggravated. Accordingly, systemic iron chelation attenuated estradiol-triggered progressive atherosclerosis in late postmenopausal mice. Thus, iron and estradiol together downregulate ERα through Mdm2-mediated proteolysis, explaining failures of HRT in late postmenopausal subjects with aging-related iron accumulation. HRT is recommended immediately after menopause along with appropriate iron chelation to protect from atherosclerosis.

## 1. Introduction

Atherosclerosis (AS) is the leading cause of cardiovascular disease-associated death worldwide (Moss *et al*, 2019). It has been well recognized that postmenopausal women confront an increasing risk of AS owing to estrogen deficiency and disturbance of the estrogen receptor (ER) regulatory network (Moss *et al*., 2019). Nevertheless, the therapeutic effect of hormone replacement therapy (HRT) remains controversial. Women’s Health Initiative and the Heart and Estrogen/Progestin Replacement Study reported that the atheroprotective effect of HRT in the late postmenopausal women (commonly ages over 65) is lost or even worsened (Hlatky *et al*, 2002; Rocca *et al*, 2014; Rossouw *et al*, 2002). The underlying mechanism may be triggered by aging, which reduces the protection afforded by estrogen, particularly estradiol (E_2_).

In general, the atheroprotective effect of estrogen is attributed to its interaction with estrogen receptor α (ERα), which participates in foam cell formation and vascular remodeling (Murphy, 2011). ERα deletion is reported to induce adiposity and increase atherosclerotic lesion size since the promoters of lipid metabolism-related genes, such as *Tgm2*, *ApoE,* and *Abca1*, contain ERα binding sites (Ribas *et al*, 2011). In addition, estrogen binds to ERα tethered to the plasma membrane, which can stimulate vasodilatation via an endothelial nitric oxide synthase (eNOS)-dependent pathway(Gavin *et al*, 2009; Teoh *et al*, 2020). Moreover, VEGF promotes angiogenesis and is transcriptionally regulated by the E_2_-ERα complex (Gu *et al*, 2018). ERα expression decreases after menopause (Gavin *et al*., 2009; Zhang *et al*, 2019), suggesting the critical role of ERα in blood vessel function in healthy pre- and postmenopausal women. Despite the above, few studies have focused on the regulation of ERα in postmenopausal women on HRT.

Upon binding, estrogen functions as an ERα activator (Lung *et al*, 2020) and initiates ERα dimerization and translocation into the nucleus. Estrogen treatment *in vitro* leads to a marked increase in *Esr1* (gene that encodes ERα) mRNA through the binding of the E_2_-ERα complex to estrogen-responsive elements (EREs) in the promoter region of *Esr1* (Pinzone *et al*, 2004). Estrogen binding to ERα also rapidly stimulates ubiquitination and proteasomal degradation of ERα (Pinzone *et al*., 2004). Ubiquitination-dependent ERα cycling on and off the ERE promoter sites to activate or prevent target gene transcription depends on the presence of estrogen at an adequate concentration (Zhou & Slingerland, 2014). Ubiquitin ligases that have been implicated in ERα regulation include BRCA1, MDM2, SKP1–CUL1–F-box S-phase kinase-associated protein 2 (SCF^SKP2^), and E6-associated protein (E6AP), all of which promote estrogen-induced transcriptional activity with cell-type selectivity (Zhou & Slingerland, 2014). Of these, MDM2 is a single-subunit RING finger E3 protein, which does not always inversely correlate with ERα levels in cancer cells (Duong *et al*, 2007). The relationship between MDM2 and ERα in other contexts and cell types remains elusive.

Estrogen also regulates systemic iron homeostasis through hepcidin, an antimicrobial peptide of 25 amino acids that mediates endocytosis and degradation of the ferrous iron exporter ferroportin 1 (Fpn1) (Nemeth *et al*, 2004). The dimeric E_2_-ERα complex binds to an ERE site within the promoter of the hepcidin gene (*Hamp*) to inhibit its expression (Hou *et al*, 2012). Hence, E_2_ elevates circulating iron to compensate for blood loss during menstruation (Badenhorst *et al*, 2021). However, E_2_ declines by over 90% after menopause, while systemic iron content increases slowly by steady iron uptake over the years. Serum ferritin in postmenopausal women increased by 2-3 times compared with premenopausal women (Huang *et al*, 2013). We and others have previously reported that iron overload is a risk factor for atherosclerosis (Cai *et al*, 2020; Vinchi *et al*, 2020). Our study relied on using the proatherogenic *ApoE^-/-^*mouse model, which we can manipulate genetically to cause iron overload (Cai *et al*, 2020). We aimed to investigate the impact of aging-related iron accumulation on the therapeutic effect of HRT on AS and explore the underlying mechanisms. First, we corroborated on a new cohort of postmenopausal AS patients that a negative correlation between high concentrations of serum-iron and -ferritin and low expression of ERα exists. We hypothesized that gradual iron accumulation in postmenopausal women could explain poor HRT efficacy when applied at a late stage.

We used *ApoE^-/-^* aging female mice or ovariectomized (OVX) young mice as estrogen-deficient AS models to examine ERα expression with respect to HRT efficacy and the genetic iron-overload mice in *ApoE^-/-^* background (*ApoE^-/-^ Fpn1^LysM/LysM^*) to address the relation of iron metabolism and HRT efficacy. We found that ERα expression positively responded to E_2_ administration under low or normal iron conditions and negatively responded to E_2_ administration under high iron conditions. Our results explain the puzzle of HRT in early and late postmenopausal women, and the reason is the accelerated ERα proteolysis through iron-mediated upregulation of *Mdm2*.

## 2. Methods and Materials

### 2.1 Participants

Participants in this study included 20 postmenopausal (at least one year since menopause, without HRT) AS patients aged 54-84 years, recruited from the Vascular Surgery Department, The Affiliated Drum Tower Hospital, Nanjing University Medical School. Patients were divided into early (55-65 years old) and late (>65 years old) groups since menopause for over ten years was defined as late post-menopause. Twenty fasting serum samples were collected at the outpatient service. Of them, eight patients, undergoing carotid endarterectomy, were recruited, and plaque samples were collected immediately after separation. Exclusion criteria included the current use of oral contraceptives or other medications. Further details on the exclusion criteria were referenced (Wactawski-Wende *et al*, 2009). This study complies with the Declaration of Helsinki, and the Institutional Review Board of Nanjing Drum Tower Hospital, the Affiliated Hospital of Nanjing University Medical School, approved the study. All patients provided written informed consent.

### 2.2 Animals

*ApoE^-/-^* mice were obtained from the Model Animal Research Center of Nanjing University (Nanjing, China). *ApoE^-/-^ Fpn1^LysM/LysM^* mice on the C57BL/6J background were generated in our previous study (Cai *et al*., 2020). For the control experiments included in Results 2 and 6, female *ApoE^-/-^* mice at the age of 8 weeks (defined as premenopausal) were anesthetized and bilaterally ovariectomized through a 1 cm dorsal incision. After surgery, mice were allowed to recover for one week and randomly divided into early and late groups. For each group, mice were fed a high-fat diet (0.2% cholesterol and 20% fat) and injected with saline, E_2_ (3 μg/kg every other day, Solarbio, Beijing, China), or E_2_+DFP (80 mg/kg daily, Sigma–Aldrich, St. Louis, MO) for eight weeks from week nine as the early OVX group or week 21 as the late OVX group.

For the control experiments included in Result 3, female *ApoE^-/-^*and *ApoE^-/-^ Fpn1^LysM/LysM^* mice were ovariectomized at the age of 16 or 40 weeks. E_2_ injection (3 μg/kg every other day) was performed one week after surgery for eight weeks. All animals were housed and fed standard chow in the SPF animal facility with an average 12 h light-and-dark cycle and under controlled temperature conditions (25°C). The mice were anesthetized with an intraperitoneal injection of pentobarbital sodium (40 mg/kg) and euthanized by cervical dislocation for sample collection. The protocols were approved by the Animal Investigation Ethics Committee of The Affiliated Drum Tower Hospital of Nanjing University Medical School and were performed according to the Guidelines for the Care and Use of Laboratory Animals published by the National Institutes of Health, USA.

### 2.3 Cell culture

J774a.1 cells, HUVECs, and MCF-7 cells were purchased from Cellcook (Guangzhou, China) and cultured in DMEM (Gibco, Life Technologies, UK) supplemented with 10% fetal bovine serum (FBS). Peritoneal macrophages were collected from peritoneal exudates three days after injecting 8-week-old mice with 0.3 ml of 4% BBL thioglycollate, Brewer modified (BD Biosciences, Shanghai, China), and then cultured in RPMI 1640 medium supplemented with 10% (FBS) for 8 h. Macrophages were cultured in a medium containing 50 μg/ml human oxidized low-density lipoprotein (oxLDL) in the presence of 1 μM E_2_, 100 μM ferric ammonium citrate (FAC), 350 μM DFP or indicated combination for 48 h as needed.

Oil Red O staining was performed to evaluate foam cell formation. The quantification was carried out with the following formula: relative average area of the fat droplet (%) = Target (Area of fat droplets/Numbers of cells)/Control (Area of fat droplets/Number of cells)*100%. The area was analyzed by ImageJ software. Angiogenesis assays were performed to evaluate angiogenic capacity. Cellular iron levels were estimated using ferrozine assays. The protein levels of ERα, ABCA1, VEGF, TfR1, and Ft-L were determined by Western blot analysis.

### 2.4 Isolation of peritoneal macrophages from mice

The mice were intraperitoneally injected with 4% starch broth (NaCl 0.5 g, beef extract 0.3 g, peptone 1.0 g, and starch 3.0 g in 100 ml of distilled H_2_O) 3 days before euthanasia. After anesthesia, the abdominal skin was carefully cut to 1 cm, and 5-8 ml PBS with 3% FBS was injected into the enterocoele. After 10 min of massage, the liquid was extracted and centrifuged (1000 rpm, 5 min). The sediment was then plated into 6-well plates for attachment or cryopreserved for further assays.

### 2.5 Blood samples and tissue collection

The mice were anesthetized with an intraperitoneal injection of pentobarbital sodium (40 mg/kg) and euthanized by cervical dislocation. Blood was drawn from the inferior vena cava and collected in heparinized tubes. Plasma was prepared by centrifugation (1200 ×g) for 15 min at 4°C. Plasma samples were then stored at -80°C for the determination of serum iron, lipid and cytokine levels. The mice were then perfused with 4°C saline through the left ventricle. After perfusion, the arteries, hearts, livers, and spleens were harvested. The samples were fixed in 4% paraformaldehyde or quickly frozen at -80°C for further analysis.

### 2.6 Serum lipid content and lesion area in the aorta and the aortic root

Lipid content was determined with Oil Red O to stain the aorta. To assess the atherosclerotic lesion area, the aorta was analyzed from the aorta arch to the abdominal aortic bifurcation. The quantification of lesion area and size was performed using ImageJ software. Serum cholesterol and triglycerides were measured by the clinical laboratory of Nanjing Drum Tower Hospital using an autochemical analyser (Beckman Coulter AU5421, CA).

### 2.7 Immunohistochemistry (IHC) and Prussian blue staining

Sections of mouse aortic valve or patient carotid artery plaques were used to assess the plaque iron composition by IHC staining for Ft-L and Prussian blue staining with DAB enhancement for ferric iron. The primary antibody against Ft-L was made using recombinant human Ft-L subunit as antigen by GenScript (Nanjing, China).

Images were captured under a light microscope (Leica, Germany). For quantitative analysis of images, three sections per animal at intervals of 30 μm were analyzed. The intensity of positive staining was analyzed by ImageJ software.

### 2.8 Iron assays

Deparaffinized tissue sections were stained with Prussian blue staining for nonheme iron as previously described (Wang *et al*, 2016). Serum iron was measured by the clinical laboratory of Nanjing Drum Tower Hospital using an autochemical analyser (Beckman Coulter AU5421, CA). Total nonheme iron in the tissues was measured by colorimetric ferrozine-based assays as previously described (Li *et al*, 2018). Briefly, 22 μl concentrated HCl (11.6 M) was added to 100 μl of homogenized tissue samples (approximately 500 μg total protein). The sample was then heated at 95°C for 20 min, followed by centrifugation at 12000 g for 10 min. The supernatant was transferred into a clean tube. Ascorbate was added to reduce the Fe3^+^ into Fe2^+^. After 2 min of incubation at room temperature, ferrozine and saturated ammonium acetate (NH4Ac) were sequentially added to each tube, and the absorbance was measured at 570 nm (BioTek ELx800, Shanghai, China) within 30 min.

### 2.9 Determination of eNOS and ferritin by ELISA

eNOS (Abcam, Cambridge, MA) and ferritin (USBiological, #F4015-11, Swampscott, MA) were detected by ELISA according to the manufacturer’s protocols.

### 2.10 Western blotting

Protein lysates were run in gels and transferred to membranes as previously reported (Cai *et al*, 2018). The membranes were probed using antibodies directed against ERα, eNOS and VEGF purchased from Servicebio (Wuhan, China), ABCA1, BRCA1, AHR, and SOD2 from Abcam, TfR1 from ProteinTech Group Inc. (Chicago, IL), GAPDH from Bioworld Tech. (St. Louis Park, MN), ferritin L (Ft-L) made by using purified human ferritin L subunit as antigen by GenScript (Nanjing, China).

### 2.11 Detection of catalase enzymatic activity

The activities of catalase were measured following the manufacturer’s protocols of the CAT assay kit (Jiancheng Bioengineering, Nanjing, China).

### 2.22 Quantitative real-time PCR (qRT–PCR)

Total cellular RNA was isolated from peritoneal macrophages using TRIzol (Invitrogen, Carlsbad, CA) and reversely transcribed to cDNA. qRT–PCR experiments were performed with SYBR Green PCR master mixture (Thermo Fisher Scientific). The primer sequences were as follows: 5’-TTATGGGGTCTGGTCCTGTG-3’ and 5’-CATCTCTCTGACGCTTGTGC-3’ for *Esr1*, 5’-GCCACTGCCGCATCCTCTTC-3’ and 5’-AGCCTCAGGGCATCGGAACC-3’ for *Actin*, and 5’-TGTCTGTGTCTACCGAGGGTG-3’ and 5’-TCCAACGGACTTTAACAACTTCA-3’ for *Mdm2*.

### 2.13 Immunoprecipitation (IP) and detection of ubiquitin

ERα proteins were immunoprecipitated from J774a.1 cell lysates according to the manufacturer’s protocols (11204D, Invitrogen). The ERα antibody was the same as that used for Western blotting and was purchased from Santa Cruz (1:200 dilution). Western blotting was used to detect the efficiency of IP and the level of ubiquitin.

### 2.14 Angiogenesis assays

Matrigel (50 μL/well) was transferred to a 96-well plate, followed by inoculation of HUVECs (2 × 10^4^ cells) and treatment with the medicines described in the cell culture. After 8 hours, images were captured with an inverted microscope. The extent of tube formation was assessed by measuring branch points and capillary length using the ‘Angiogenesis Analyser’ plug-in designed by Gilles Carpentier with ImageJ software.

### 2.15 Statistical analysis

All experiments were randomized and blinded. All the data are presented as the mean ± SD. A two-tailed Student’s t-test (for two groups) or one-way analysis of variance followed by multiple comparisons test with Bonferroni correction (for more than two groups) was performed by using SPSS 17.0 (SPSS Inc, Chicago, IL). P < 0.05 indicated statistical significance.

## 3. Results

### 3.1 ERα protein abundance correlates inversely with systemic and local plaque iron content in aging postmenopausal women with AS

To build the link between iron contents and ERα levels in different postmenopausal stages, we collected 8 plaques and 20 blood samples from postmenopausal AS patients, divided a half into Early (55-65 years old, n=4/10, plaque/blood samples) and another half into Late (>66 years old, n=4/10) Post-Menopausal groups, EPM and LPM, respectively. Ferritin was evaluated by IHC, ELISA, and immunoblotting, and iron was evaluated by DAB-enhanced Prussian blue staining. As expected, both ferritin and iron levels were significantly increased in plaques and serum of the LPM, compared to the EPM (Figure 1A-C, E). In addition, serum cholesterol and triglycerides were elevated in the LPM (Figure 1D). By contrast, plaque ERα expression was lower in the EPM than in the LPM (Figure 1E).

**Figure 1.**
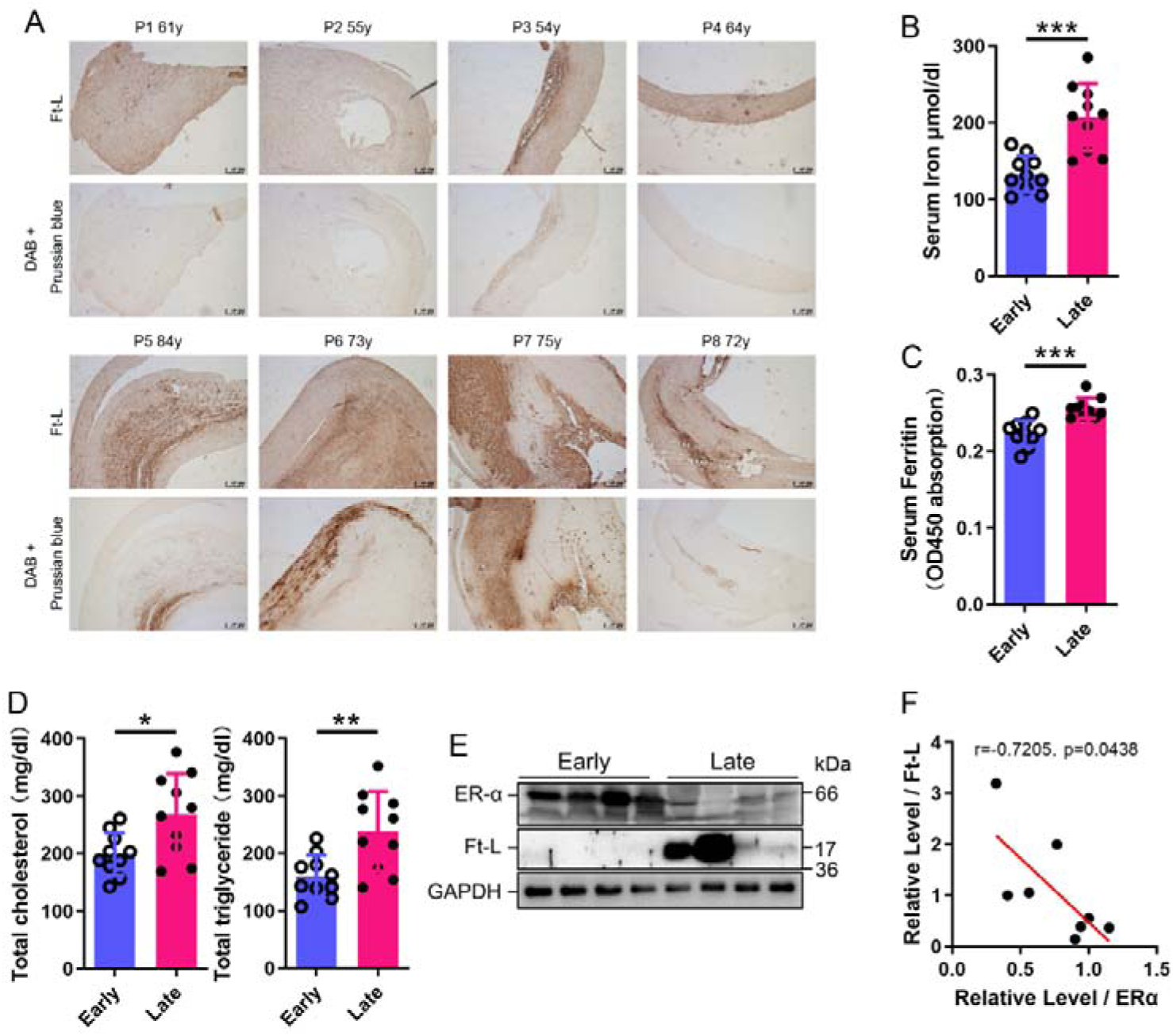
ERα levels were negatively associated with iron content in human plaques. (A) Ferritin (Ft-L), revealed by immunohistochemistry, and iron content, revealed by DAB-enhanced Prussian blue staining, in plaque paraffin sections of 8 postmenopausal patients. The upper panel: the early postmenopausal (EPM) group (P1-P4, < 65 years old); the lower panel: the late postmenopausal (LPM) group (P5-P8, > 65 years old). (B) Serum iron in EPM (blue) and LPM (magenta) patients, measured by using an autochemical analyser (Beckman Coulter AU5421, CA). n=10/group, \*\*\**p* < 0.001. (C) Serum ferritin levels detected by ELISA. n=10/group, \*\*\**p* < 0.001. (D) Serum total cholesterol (left) and total triglyceride (right) levels. n=10/group, \**p* < 0.05, ** *p* < 0.01. (E) ERα and Ft-L expression in plaques measured by Western blotting. The samples are the same as in (A). (F) The plotted and calculated Pearson correlation coefficient (*r =* -0.7205) between the plaque Ft-L and ERα levels (n=8, *p* = 0.0438).

Pearson correlation analysis between ERα and ferritin levels in the plaques confirmed that ERα levels were negatively correlated with iron levels (Figure 1F).

### 3.2 AS aggravates in E_2_-treated LPM *ApoE^-/-^* mice with reduced ERα expression and accumulation of body iron

To determine whether the effect of HRT was atheroprotective in postmenopausal females, ovariectomy (OVX) was performed to mimic post-menopause in *ApoE^-/-^* female mice at eight weeks of age (Figure 2A). The mice started to be fed high-fat chow one week post OVX. Ages of 9 weeks and 21 weeks were considered as EPM and LPM stages, respectively. Although peritoneal E_2_ injection in the EPM significantly reduced plaque formation, it remarkably promoted atherosclerotic development in the late application (Figure 2B and 2C), exactly as observed in humans (Hlatky *et al*., 2002; Rossouw *et al*., 2002). We then detected aortic ERα expression to evaluate whether ERα was responsive to E_2_ treatment. The results showed markedly lower ERα protein levels in the LPM mice than in the EPM mice (Figure 2D). More strikingly, ERα expression was further reduced in LPM mice after E_2_ treatment but remained constantly high in EPM mice (Figure 2D). It has been reported that ERα protects against atherosclerosis by promoting lipid efflux and endothelial homeostasis (Wang *et al*, 2021; Zhao *et al*, 2021). Hence, we assessed three ERα downstream proteins, ABCA1, a lipid exporter whose gene promoter is predicted to have ERE, VEGF, an activator of angiogenesis, and eNOS, a modulator of vasoconstriction and vascular repair. They were all positively correlated with ERα expression (Figure 2D). Macrophage-derived foam cell formation is crucial in the development of atherogenesis (Xu *et al*, 2021). We therefore isolated peritoneal macrophages from early and late OVX mice after E2 treatment and surprisingly found that the expression of ERα, ABCA1, and VEGF responded to E_2_ treatment similarly as observed in aortic tissue (Figure 2E). In line with this observation, serum cholesterol and triglycerides negatively correlated with ERα and ABCA1 (Figure 2F). Our previous data have demonstrated that macrophage iron plays a critical role in the development of AS (Cai *et al*., 2020); therefore, iron-related proteins were monitored.

**Figure 2.**
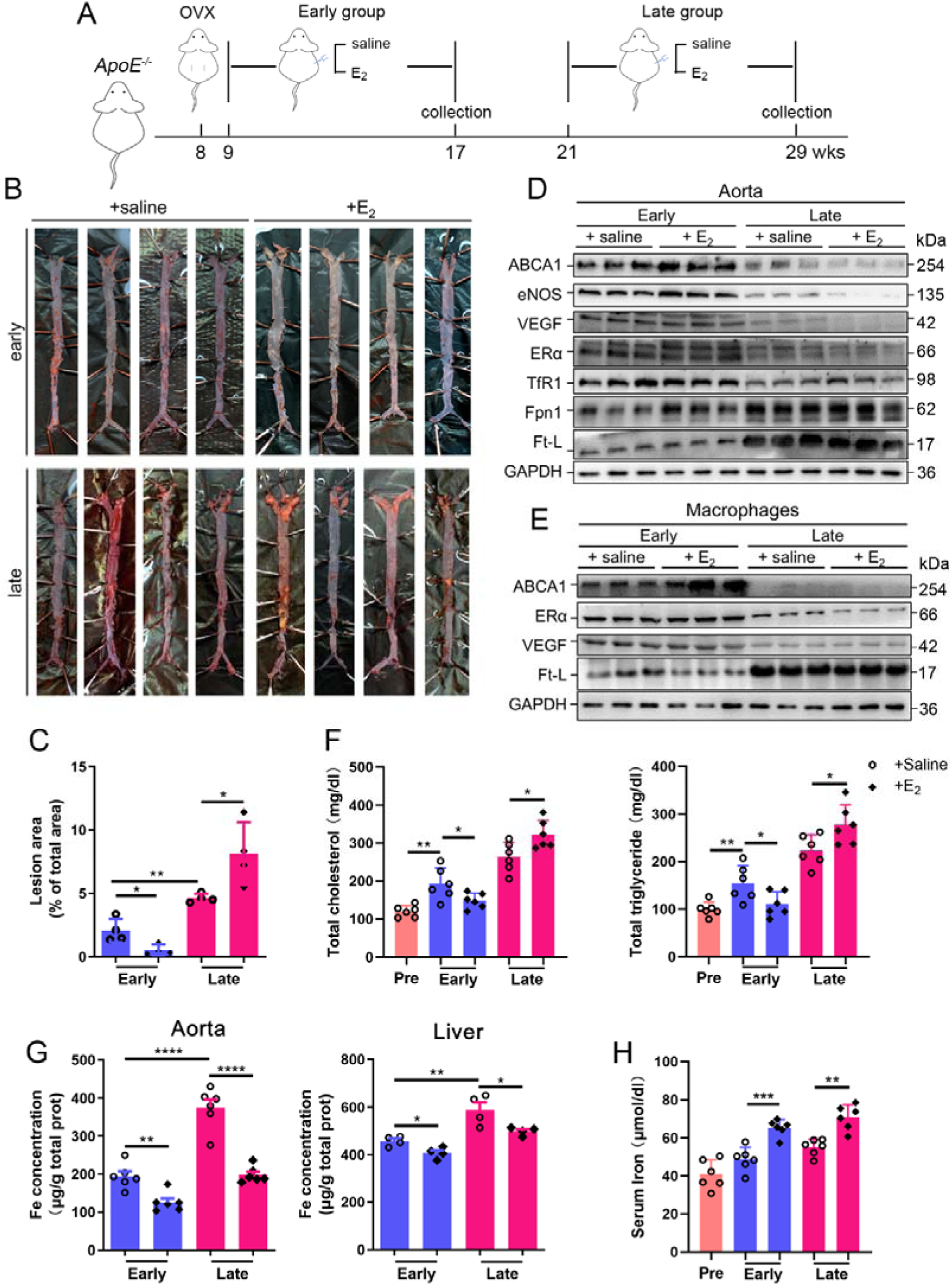
Atherosclerosis was aggravated in E_2_-treated late postmenopausal *ApoE^-/-^* mice with lower ERα expression. (A) Flow diagram of mouse modeling. Early E_2_-treatment group: OVX at 8 weeks old, one-week recovery, E_2_ treatment for 8 weeks; late E_2_-treatment group: OVX at 8 weeks old, E_2_ treatment from 21 weeks old to 29 weeks old for 8 weeks. Saline is vehicle control. Mice were fed high-fat chow from 9 weeks old. (B) Oil red O-stained aortic lesions in *ApoE^-/-^* mice after E_2_ treatment for 8 weeks in the EPM or LPM group. (C) Statistical analysis of the area of atherosclerotic plaque in the aorta. n = 4/group, \**p* < 0.05, ** *p* < 0.01. (D) The expression of iron-related or ERα-targeted proteins in the aorta, detected by Western blotting. (E) Protein expression in peritoneal macrophages detected by Western blotting. Macrophages were isolated from 4 mouse groups (early/late ± E2, details see **Materials and Methods**). (F) Serum total cholesterol and total triglyceride levels in the 4 mouse groups. Pre: serum samples before OVX as a control group. n=6/group, \**p* < 0.05, ** *p* < 0.01. (G) Iron content in aorta and liver, detected by ferrozine assays. n = 6/group, \*\*\*\**p* < 0.0001, \*\**p* < 0.01, \**p* < 0.05. (H) Serum iron in different groups, detected by using an autochemical analyser (Beckman Coulter AU5421). n = 6/group, \*\*\**p* < 0.001, \*\**p* < 0.01. Student’s *t-test* analysis was used for C, F, G, and H.

Ferritin was reduced in response to E_2_ treatment in the EPM stage. In contrast, ferritin remained high in the late stage after E_2_ treatment in both aortae and isolated macrophages (Figure 2D and 2E), which could be explained by the response of Fpn1 expression that was decreased in the EPM stage but not in the LPM stage. Both serum and tissue iron levels were significantly higher in the LPM mice (Figure 2G-2H, LPM vs. EPM without E_2_ treatment). Interestingly, E_2_ treatment elevated serum iron while lowering tissue iron in both EPM and LPM stages (Figure 2G-H, E_2_ treatment vs. saline) suggesting impaired iron homeostasis in the plaque area, particularly in macrophages (Figure 2E) and confirming that estrogen modulates iron homeostasis as previously suggested (Yang *et al*, 2012).

### 3.3 E_2_ downregulates ERα expression in an iron-dependent manner

Next, we aimed to identify whether aging or iron overload alone could trigger a decrease in ERα expression *in vivo*. To address this question, myeloid-specific *Fpn1* knockout mice (*Fpn1^LysM/LysM^*) were used as a macrophage-iron overload model in the *ApoE^-/-^* background (Cai *et al*., 2020). This double knockout (KO) model is considered relevant for AS studies due to the accumulation of a large number of macrophages in plaques, which contributes to the progression of atherosclerosis (Moore *et al*, 2013). OVX was performed in female mice fed standard chow at 16 weeks or 40 weeks of age, and E_2_ was injected to model HRT, as illustrated in Figure 3A. Figures 3B and 3C show the severity of atherosclerosis, which was significantly enhanced in the E_2_-treated groups at both ages compared to the saline groups of *ApoE^-/-^ Fpn1^LysM/LysM^*. Notably, macrophage *Fpn1* KO mice displayed a larger lesion area after E_2_ treatment at the EPM and LPM stages (Figure 3C), suggesting a dominant influence of iron on the effects of the E_2_ treatment. In particular, the specific iron overload in macrophages, characteristic of the mouse model used, was sufficient to cause a significant increase in the lesion area in the LPM group (Figure 3C, lower panel), reproducing previous observations (Cai *et al*., 2020). We then examined the iron status in tissues and serum. Iron levels in tissues (aorta/liver) were higher in *ApoE^-/-^ Fpn1^LysM/LysM^* compared to *ApoE^-/-^* mice, as revealed by ferrozine assays (Figure S1A for EPM groups and Figure 3E for LPM groups) and further supported by higher ferritin content (Figure S1B for EPM and Figure 3D for LPM). On the contrary, serum iron was lower in *ApoE^-/-^Fpn1^LysM/LysM^* mice at the EPM and LPM stages (Figure 3F and S1C). However, E_2_ administration did not significantly reduce the ferritin levels and iron levels in the aorta and liver (Figure 3D, 3E, and Figure S1A), confirming that Fpn1 acts as an iron exporter and that macrophages play a crucial role in response to E_2_ treatment. E_2_ administration significantly increased serum iron levels in the LPM group (Figure 3F) and mildly increased serum iron levels in the EPM group (Figure S1C), suggesting that other factors besides iron also contribute to the aging-related physiological changes in the E_2_ treatment response. Of note, ERα was downregulated in the aorta of the *Fpn1^LysM/LysM^*mice and further downregulated by E_2_ treatment, accompanied by reduced expression of ABCA1 and VEGF (Figure 3B and Figure S1B), suggesting that both iron alone and iron plus E_2_ could downregulate ERα expression in macrophages. Consistent with the severity of atherosclerosis and the alteration of ERα and its target gene ABCA1, serum cholesterol and triglycerides were increased in the *Fpn1^LysM/LysM^*mice and further increased by E_2_ treatment (Figure 3G).

**Figure 3.**
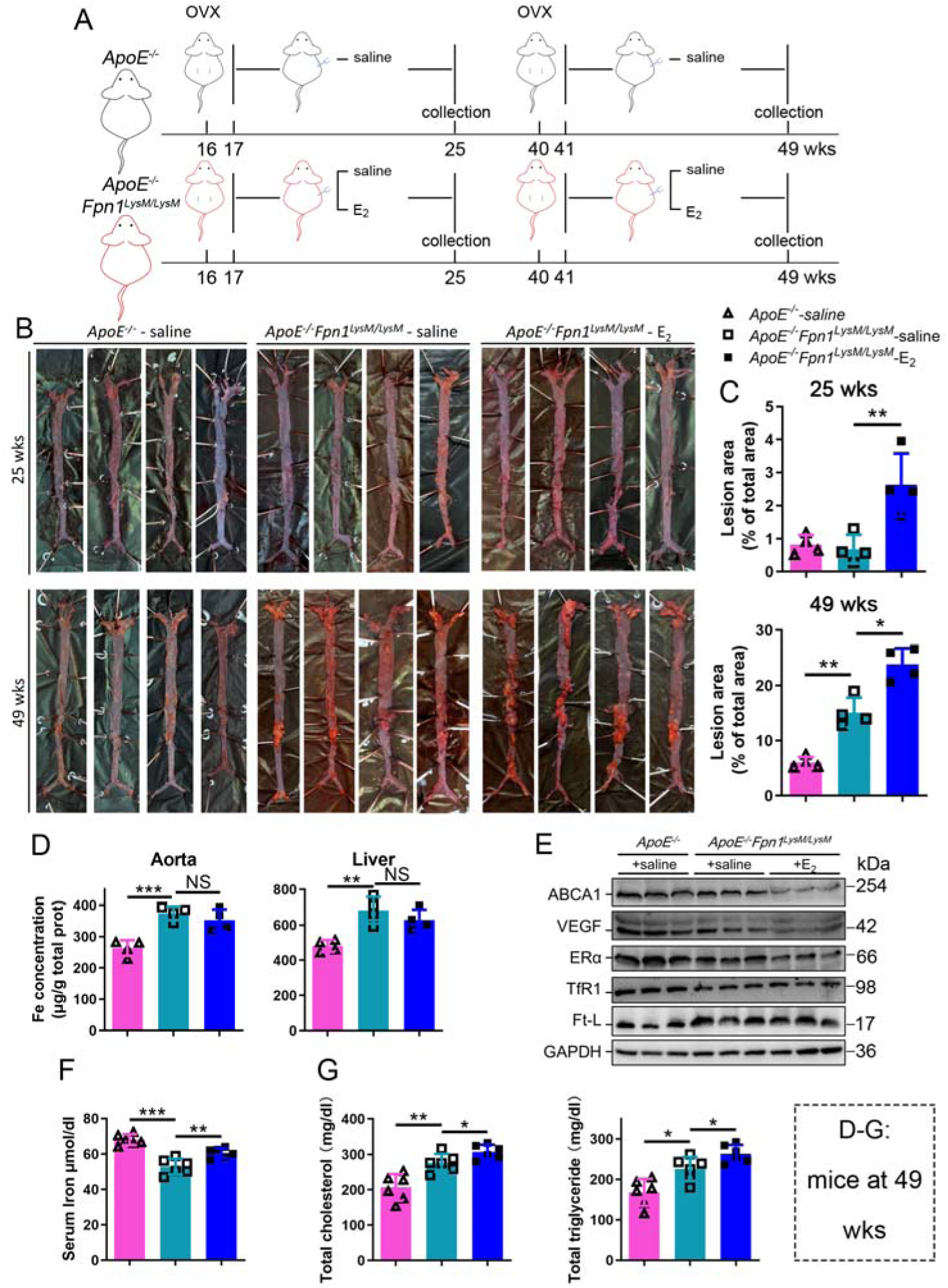
E_2_-triggered ERα deficiency was observed in a genetic iron overload mouse model at postmenopausal age. (A) Flow diagram of mouse modeling. Early groups: OVX at 16 weeks old, one-week recovery, ± E_2_ treatment for 8 weeks; late groups: OVX at 40 weeks old, one-week recovery, ± E_2_ treatment for 8 weeks. Saline is vehicle control. The mice were fed with normal chow. (B) Oil red O-stained aortic lesions in *ApoE^-/-^*and *ApoE^-/-^ Fpn1^LysM/LysM^* mice after E2 treatment for 8 weeks in the EPM or LPM groups as indicated. (C) The lesion area in the aorta. n = 4/group, \*\**p* < 0.01, \**p* < 0.05. (D) The expression of iron-related or ERα-targeted proteins in the aorta, detected by Western blotting. (E) The iron content of the aorta and liver detected by ferrozine assays. n = 6/group, \*\*\**p* < 0.001, \*\**p* < 0.01. (F) Serum iron level in different groups. n = 6/group, \*\*\**p* < 0.001, \*\**p* < 0.01. (G) Serum total cholesterol and total triglyceride levels. n=6/group, \**p* < 0.05, ** *p* < 0.01. The samples for D-G were from 49-week-old *ApoE^-/-^* and *ApoE^-/-^ Fpn1^LysM/LysM^* mice. Student’s *t-test* analysis was used for C, E, F, and G.

### 3.4 E_2_ treatment potentiates iron-induced downregulation of ERα in both macrophages and endothelial cells

To further validate the interaction between iron and E_2_ on ERα downregulation in different cell types, we used primary peritoneal macrophages from C57BL/6 female mice, the macrophage-like cell line J774a.1, and human umbilical vein endothelial cells (HUVECs). The cells were treated with E_2_, ferric ammonium citrate (FAC, an iron source), and/or deferiprone (DFP, an iron chelator). Downregulation of ERα expression triggered by FAC with or without E_2_ was observed in time- and concentration-dependent manners (Figure 4A and 4B). Such downregulation was confirmed in all tested cell types and could be partially suppressed by iron chelation (Figure 4C, 4D, 4E, and 4F).

**Figure 4.**
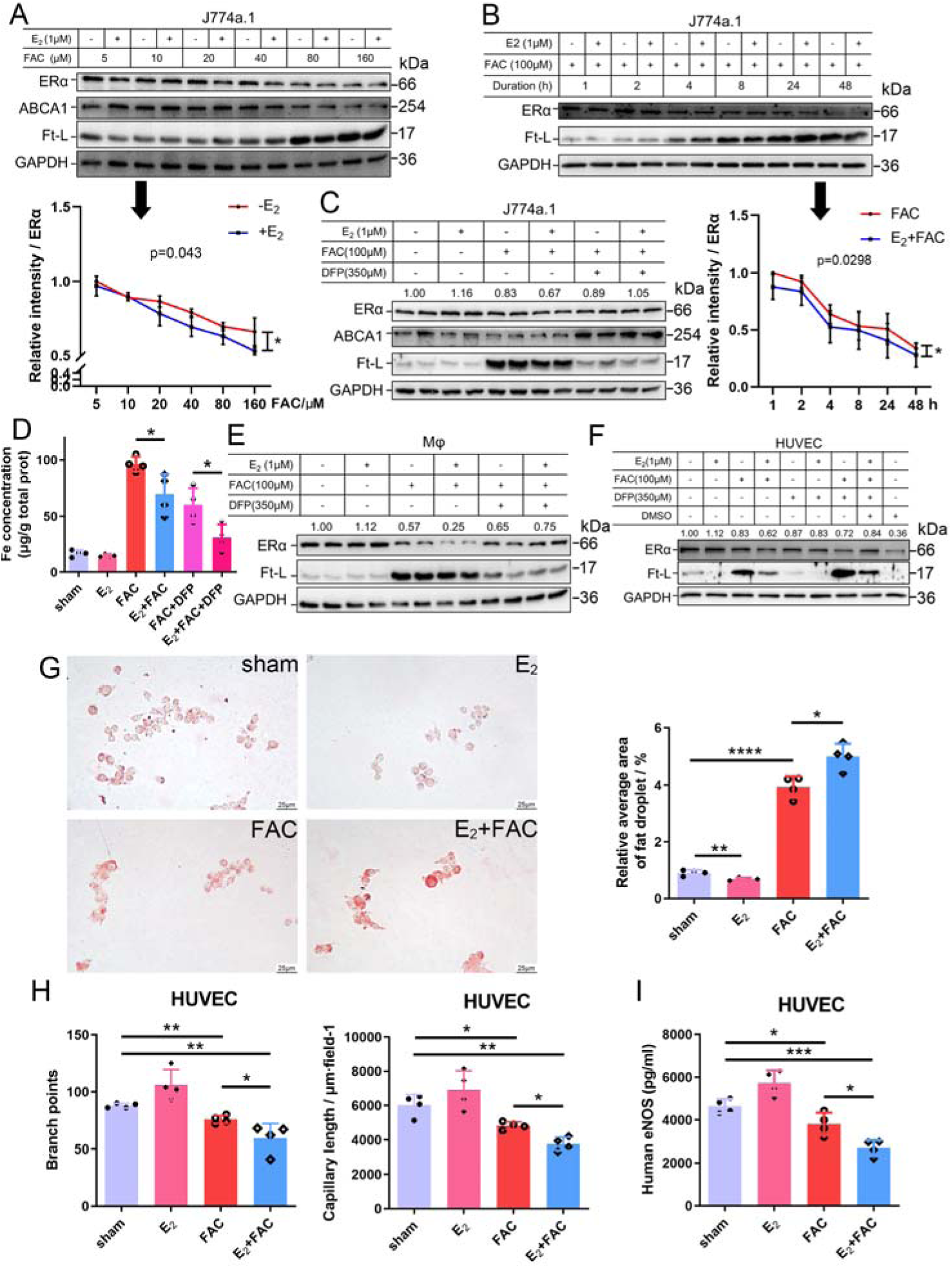
E_2_ treatment potentiates iron-induced downregulation of ERα *in vitro*. (A-B) ERα expression in the presence or absence of E_2_ under different iron concentration conditions (A) or in the time course (B). Quantification was carried out using ImageJ. Two-way ANOVA was used. (C) The rescue effect of iron chelation on the downregulation of ERα by FAC or FAC plus E_2_. (D) The intracellular iron content in J774a.1 under different iron-concentration conditions in the presence or absence of E_2_, detected by ferrozine assays. n = 4, \**p* < 0.05. (E, F) ERα expression in peritoneal macrophages (E) and HUVECs (F) under the indicated iron and E2 conditions. A, B, C, E, and F are data from Western blotting. Numbers indicate the relative intensity of ERα n=4. (G) Oil red O-stained J774a.1 cells after treatment with FAC and/or E_2_ (left) with the quantified droplets (right). scale bar = 25 μm, n = 4, \*\*\**p* < 0.001. (H) HUVEC angiogenesis assays, revealed by the number of branch points (left) and capillary length (right). n = 4, \**p* < 0.05, \*\**p* < 0.01. (I) eNOS level in HUVEC, assessed by ELISA. n = 4, \**p* < 0.05, \*\**p* < 0.01, \*\*\**p* < 0.001. The student’s *t-test* analysis was used for G - I.

To examine the capacity of lipid export when ERα was downregulated, we loaded J774a.1 with oxidized low-density lipoprotein (oxLDL) and observed significantly more lipid accumulation in the E_2_-treated plus iron overload group than in the other groups (Figure 4G), suggesting a tendency of macrophages to be converted into foam cells. Angiogenesis assays were also performed and showed that E_2,_ together with iron, inhibited angiogenesis (Figure 4H), which has been demonstrated to increase the risk of macrophage adhesion and intraplaque hemorrhage (Chang & Nguyen, 2021; Mao *et al*, 2020). The reduced levels of eNOS were also revealed by ELISA in HUVECs treated with E_2_ and iron (Figure 4I). Overall, our data strongly support that both macrophages and endothelial cells are the effectors of E_2_ in iron-mediated worsening by downregulation of ERα in the development of AS.

### 3.5 Proteasome-mediated ERα degradation results from the interactive effects of iron overload and E_2_ treatment mediated by the E3 ligase Mdm2

We wondered how excess iron and E_2_ together downregulated ERα expression. To elucidate the underlying mechanism, we first searched for what regulates ERα expression. It was reported that the downregulation of ERα could be attributed to methylation of its promoter region, which could be induced by oxidative stress (Lung *et al*., 2020). Because iron overload has been massively correlated with oxidative stress, two relevant antioxidative enzymes, catalase (CAT) and superoxide dismutase (SOD) 2, were evaluated. The results did not show a significant difference between the control and E_2_/FAC treatments (Figure S2A and S2B). In addition, the mRNA level of ERα was examined and barely showed significant changes after FAC and E_2_ treatment (Figure S2C), suggesting posttranscriptional regulation of ERα better explained the decreased presence of the receptor after iron and/or E_2_ treatment.

As stated previously, ERα may be regulated through an estrogen-ERα binding-dependent ubiquitination signaling pathway for degradation and estrogen recycling. To test this possibility, we treated J774a.1 cells with MG132, a proteasome inhibitor, and observed that ERα protein levels were significantly elevated in the presence of E_2_ and excess iron (Figure 5A). Treatment with cycloheximide (CHX), an inhibitor of eukaryotic translation, showed that the half-life of ERα was shortened in the E_2_ + FAC group (Figure 5B), suggesting a faster turnover rate of ERα in the presence of E_2_ plus excess iron, verifying the activation of ERα proteasome degradation pathway (Zhou & Slingerland, 2014). We then detected the ubiquitination levels of ERα by immunoprecipitation and immunoblotting. The results showed much more ubiquitinated and degraded ERα in the presence of E_2_ and excess iron than in other conditions (Figure 5C), further supporting the proteolysis-dependent pathway for ERα degradation. We then tested a few E3 ligases (BRCA1, AHR, and Mdm2) in J774a.1, which were supposed to regulate ERα (Fan *et al*, 2001; Khan *et al*, 2006; Saji *et al*, 2001), and did not find a negative correlation between BRCA1/AHR and ERα in response to FAC or E_2_+FAC treatments (Figure S3A). However, *Mdm2* was upregulated in the FAC and E_2_+FAC groups (Figure 5D). When cells J774a.1 and HUVECs were treated with Nutlin-3, a well-known Mdm2 inhibitor, ERα exhibited iron-dependent Mdm2-mediated degradation (Figure 5E and 5F). Amazingly and consistently, Mdm2 expression was upregulated at the LPM stage compared with the EPM stage both in the aorta of female mice and in plaques of AS patients (Figures 5G and 5H), which is exactly opposite to ERα expression (Figure 1E). And this effect was significantly enhanced at the LPM stage mice when E2 was administrated (Figure 5G, 2D and 2E). Our results indicate that Mdm2 is responsible for E_2_-triggered ERα deficiency under iron overload condition or in LPM women.

**Figure 5.**
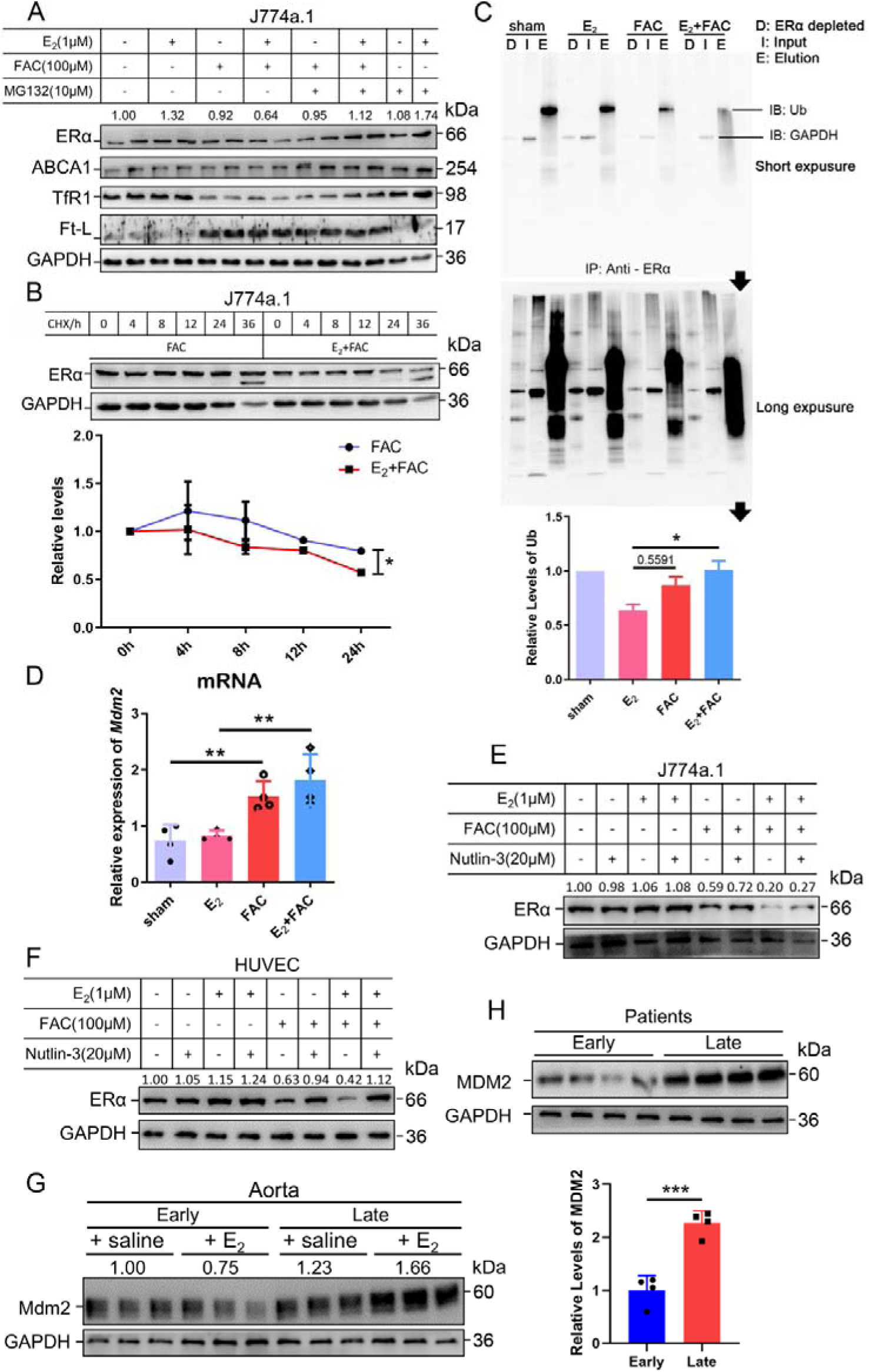
The interactive effects of iron overload and E_2_ treatment on ERα downregulation are mediated by the E3 ligase MDM2. (A) Evaluation of ERα proteasome-dependent degradation in J774a.1 cells by western blotting. MG132: 10 μM. n=4. (B) ERα turnover rate in J774a.1 cells under FAC or E_2_+FAC conditions, detected by western blotting after 20 μM cycloheximide (CHX) treatment. \**p* < 0.05 using two-way ANOVA (C) Ubiquitination of ERα, evaluated by western blotting (anti-ubiquitin) following immunoprecipitation against ERα antibody. n = 3, \**p* < 0.05. (D) Relative *Mdm*2 mRNA expression in J774a.1 cells, assessed by qPCR, n = 4, \*\**p* < 0.01. (E) The protein levels of ERα in the presence of FAC or FAC plus E_2_ in J774a.1 cells after treatment of Nutlin-3, a specific antagonist of Mdm2. n=3. (F) The protein levels of ERα in the presence of FAC or FAC plus E_2_ in HUVECs after treatment of Nutlin-3. n=3. (G) Mdm2 protein expression in the aortas of mice in the EPM or LPM stage, as detected by western blotting. n=3/group. (H) MDM2 protein levels in patient plaques, detected by western blotting and quantified with ImageJ. n=4/group, \*\*\**p* < 0.001. Student’s *t-test* analysis was used for B, E, and H.

### 3.6 Iron restriction therapy restores ERα levels and attenuates E2-triggered progressive atherosclerosis in late postmenopausal mice

To further verify whether iron overload is responsible for the E_2_-induced downregulation of ERα and progressive atherosclerosis in LPM mice, we evaluated the effects of iron restriction. Twenty-one-week-old female *ApoE^-/-^*mice, OVX-ed at eight weeks of age, received iron chelation therapy through peritoneal injection of DFP (80 mg/kg) daily for eight weeks (Figure 6A). Similar to the definition used in Figure 2, 13 weeks after OVX was considered as the LPM stage. Indeed, iron chelation attenuated the plaque-accelerated development of AS (Figure 6B and 6C). The contents of serum cholesterol and triglycerides compared with those in the E_2_-only group were significantly diminished (Figure 6D). Consistent with previous data (Figures 2 and 3), ferrozine assays proved decreased iron deposition in tissues but increased iron in serum post E_2_ application (Figure 6E-F). Although DFP administration reduced the tissue and serum iron levels, it did not induce anemia (Figure 6E-F), which was comparable to the mice at EPM (Figure 2G and H). We then detected aortic ERα expression and found significant upregulation by iron restriction, along with the upregulation of ABCA1 and VEGF (Figure 6G). In contrast, iron chelation by DFP significantly reduced Mdm2 expression (Figure 6G). In agreement with our previous findings in cell-based assays, these results corroborate the concept that late postmenopausal HRT-induced ERα deficiency is, at least partially, iron overload-mediated. Thus, the non-atheroprotective effects of E_2_ in the LPM result from aging-mediated iron accumulation.

**Figure 6.**
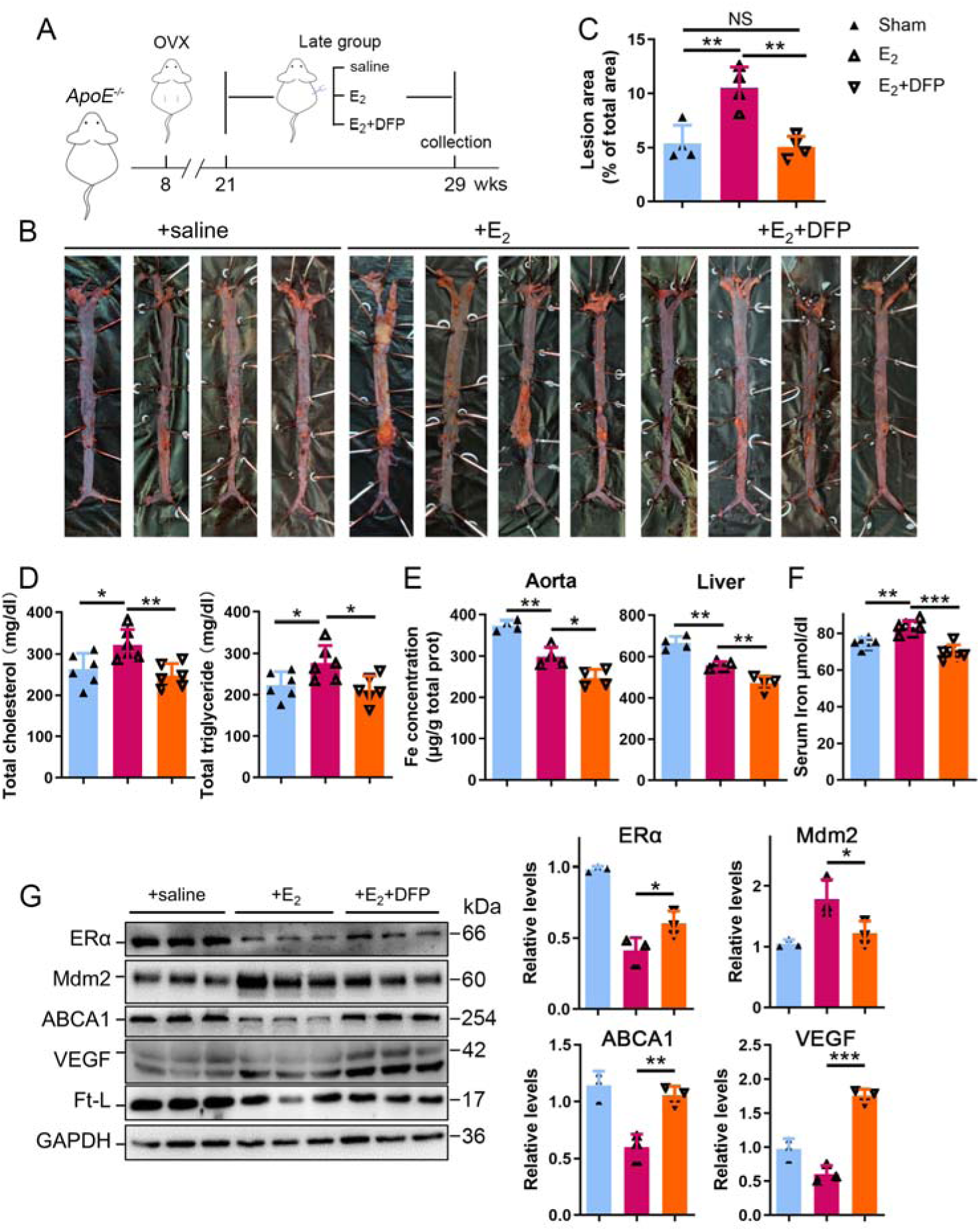
Iron restriction therapy restored ERα levels and attenuated E_2_-triggered progressive atherosclerosis in late postmenopausal mice. (A) Flow diagram of mouse modeling. The mice were ovariectomized at 8 weeks old and E_2_ or E_2_+DFP treated from 21 weeks old to 29 weeks old for 8 weeks. Saline is vehicle control. Mice were fed high-fat chow one week after OVX. Thirteen weeks post-OVX is considered as late post-menopause. (B) Oil red O-stained aortic lesions in *ApoE^-/-^* mice treated with E_2_ or E_2_+DFP as indicated. (C) The quantified lesion area of atherosclerotic plaques in the aorta from B. n=4, \*\**p* < 0.01. (D) Serum total cholesterol and total triglyceride levels. n=6, \**p* < 0.05, ** *p* < 0.01. (E) The iron content in the aorta and liver, detected by ferrozine assays. n = 6, \*\**p* < 0.01, \**p* < 0.05. (F) Determination of serum iron in different groups. n = 6, \*\*\**p* < 0.001, \*\**p* < 0.01. (G) Protein expression in the aorta, detected by western blotting (left) and quantified with ImageJ (right). \*\*\**p* < 0.001, \*\**p* < 0.01, \**p* < 0.05. The student’s *t-*test analysis was used for C - G.

## 4. Discussion

Estrogen has long been considered atheroprotective and responsible for the low morbidity of cardiovascular diseases in premenopausal women (Lobo, 2017; Moss *et al*., 2019). However, epidemiological studies of the Women’s Health Initiative question the beneficial effects of late postmenopausal HRT (Hlatky *et al*., 2002). Although one hypothesis to explain this observation may be that iron potentiates the adverse effects of estrogen in AS (Sullivan, 1981; Sullivan, 2003), a comprehensive *in vivo* study to test this hypothesis was missing. We reported previously that the developmental course of atherosclerosis was highly accelerated in *ApoE^-/-^ Fpn1^LysM/LysM^* mice compared with *ApoE^-/-^*(Cai *et al*., 2020). Our present study provides the first experimental evidence that iron overload facilitates ERα proteolysis, which is potentiated in the presence of E_2_ and reverses the anti-atherogenic effect of E_2_ (Figure 7). Our results support the benefit of early application of estrogen post-menopause. We propose that the combination of HRT and iron restriction therapy may be a long-term strategy for the preventive effects of E_2_ from the development of AS in post-menopausal women.

**Figure 7.**
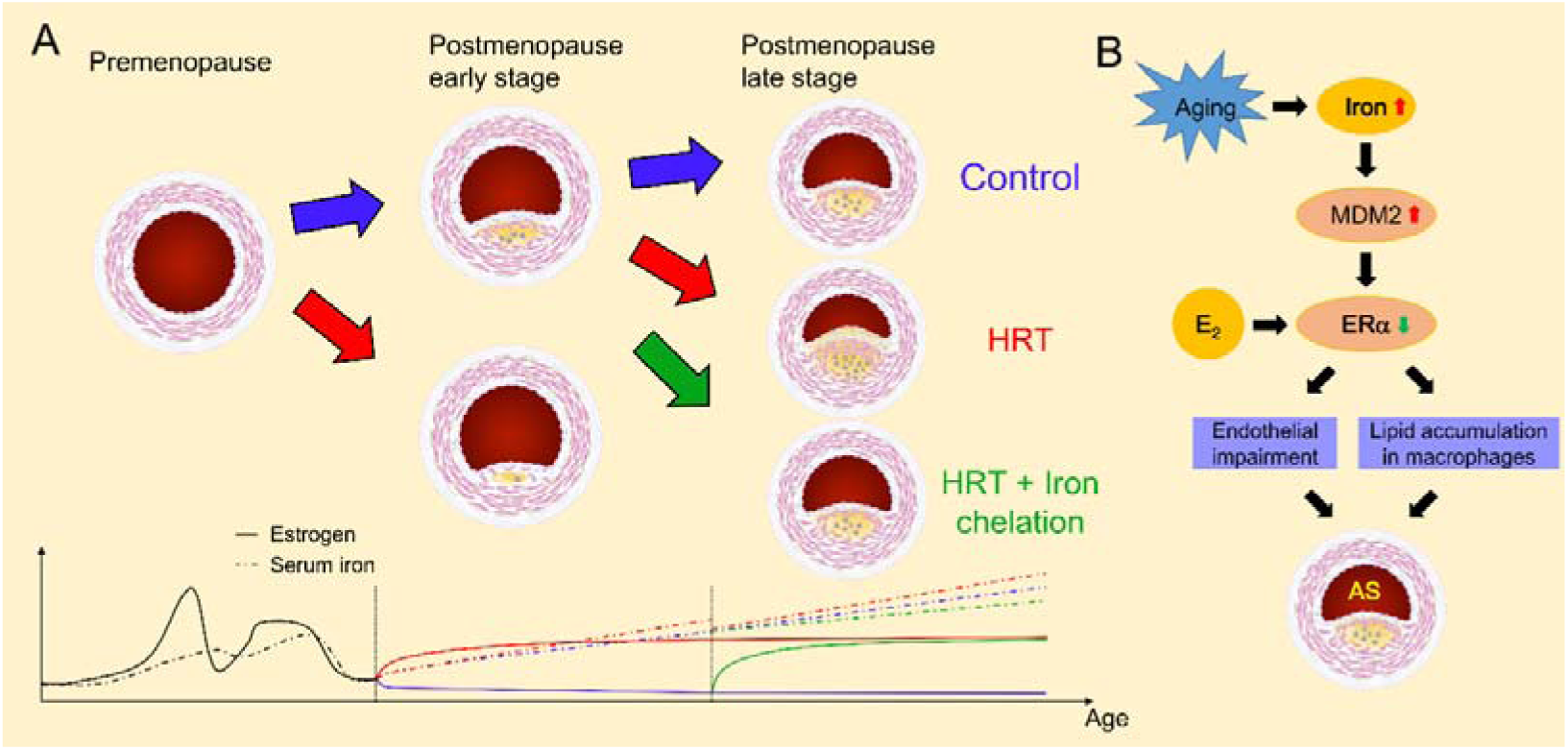
Schematic model for the effects of postmenopausal iron accumulation with or without HRT on AS severity through modulating ERα expression. Iron accumulation occurs naturally and gradually after menopause. In EPM, iron retention was mild, and ERα was responsive to HRT application to achieve protective effects (A). However, when iron overload is significant in LPM, Mdm2 is upregulated along with ERα downregulation (B). This negative correlation is potentiated by the application of HRT and iron accumulation with aging. Therefore, HRT use avails to aggravate the progression of AS in the LPM period. Iron chelation, however, reverses the adverse effect of HRT and attenuates the accelerated development of AS (A), suggesting a protective role of appropriate iron restriction in the LPM stage.

The controversy of whether or not to proceed with HRT in postmenopausal women is fueled by an increased, although small, risk of breast cancer and from the potentially harmful effect on cardiovascular outcomes (Lobo, 2017). Previous randomized trials did not consider the ages sorting out EPM from LPM and have not excluded subjects with iron depletion or loss in the recruited post-menopausal subjects (Sullivan, 2003). We sorted the recruited female AS volunteers from the Department of Vascular Surgery of Nanjing Drum Tower Hospital as EPM and LPM groups to reveal whether aging-associated iron deposition correlates with ERα expression. The negative correlation between systemic iron status and intraplaque ERα expression was validated, which prompted us to address the role of iron in ERα expression.

Previous efforts focused more on the role of estrogen in iron metabolism (primarily the hepcidin/Fpn axis) and not *vice versa* (Hou *et al*., 2012; Ikeda *et al*, 2012; Yang *et al*., 2012). Both genes encoding hepcidin and Fpn are inhibited by E_2_ treatment through an estrogen-responsive element (ERE) (Hou *et al*., 2012; Qian *et al*, 2015; Yang *et al*., 2012). However, it was also reported that hepcidin expression decreased in the livers of OVX mice through a GPR30-BMP6-dependent mechanism, independent of the ERE-mediated E_2_-ERα pathway (Ikeda *et al*., 2012). Though the difference in hepcidin expression in OVX mice (Bowling *et al*, 2014; Gavin *et al*., 2009; Hou *et al*., 2012; Ikeda *et al*., 2012), the consistent with this study here is that aging, OVX, and genetic manipulation of Fpn induced progressive iron retention in tissues, accompanied by reduced ERα expression. E_2_ administration further enhanced this reduction in ERα levels under the above-mentioned conditions. Overall, high ERα levels are found in reproductive women, despite fluctuations caused by the periodic estrogen wave and blood loss in reproductive women (Gavin *et al*., 2009). The aging process, particularly in late postmenopausal women, progressively elevates iron levels which we show, downregulates ERα, resulting in insufficient ERα to respond to E_2_ treatment. Therefore, HRT is unlikely to result in an effective outcome in LPM women as in EPM women unless it is coupled with an iron-chelating scheme. This is because aggravated AS in LPM women is, at least partially, the result of age-related iron accumulation. We demonstrated the effectiveness of iron chelation in improving HRT outcomes in the mouse model, but further work is required to translate this finding for clinical practice.

ERα is the main effector of estrogen on cardiovascular function (Aryan *et al*, 2020; Meng *et al*, 2021). We wondered how iron downregulated ERα. Since several E3 ubiquitin ligases (*i.e*., CHIP, E6AP, BRCA1, BARD1, SKP2, and Mdm2) have been found to catalyze the covalent binding of ubiquitin to lysine residues of ERα (reviewed in (Tecalco-Cruz & Ramirez-Jarquin, 2017), we tested them and found that Mdm2 is responsive to iron treatment in cells and mice and negatively correlated with ERα, particularly under high iron conditions. Furthermore, we showed that Mdm2 is a negative regulator of ERα.

Our findings may be context specific, as some differences are noted in its studied roles in some cancer cell types (Dongiovanni *et al*, 2010; Zhang *et al*, 2020). Mdm2 acts as a ubiquitin ligase E3 to p53 in SV40 hepatocytes (Honda *et al*, 1997) and has been shown to act as a direct coactivator of ERα function in ERα-positive breast cancer (Saji *et al*., 2001). Nevertheless, iron-dependent downregulation was revealed in leukemia cell lines and primary human cells derived from acute myeloid leukemia patients (Calabrese *et al*, 2020), suggesting a cell-type-specific regulation of Mdm2 by iron. In our study, the downregulation of Mdm2 by E_2_ occurred in the context of iron overload both *in vivo* and *in vitro*, concluding that Mdm2 is the critical mediator that participates in iron overload triggered ERα loss.

In summary, this study demonstrates the impact of iron overload in E_2_-mediated ERα proteolysis and its critical consequence on the outcome of HRT. With the efficacy of HRT challenged by “the window of opportunity” theory (Yesufu *et al*, 2007), it is vital to explore therapies that maintain ERα expression mediating the protective effects of estrogen. Although further work is needed to determine whether iron restriction therapy is clinically relevant in combination with HRT for the intervention of postmenopausal atherosclerosis as a long-term strategy, this paper provides guidance for optimizing the timing HRT intervention and supportive nutrient management.

---

**Figure S1.**
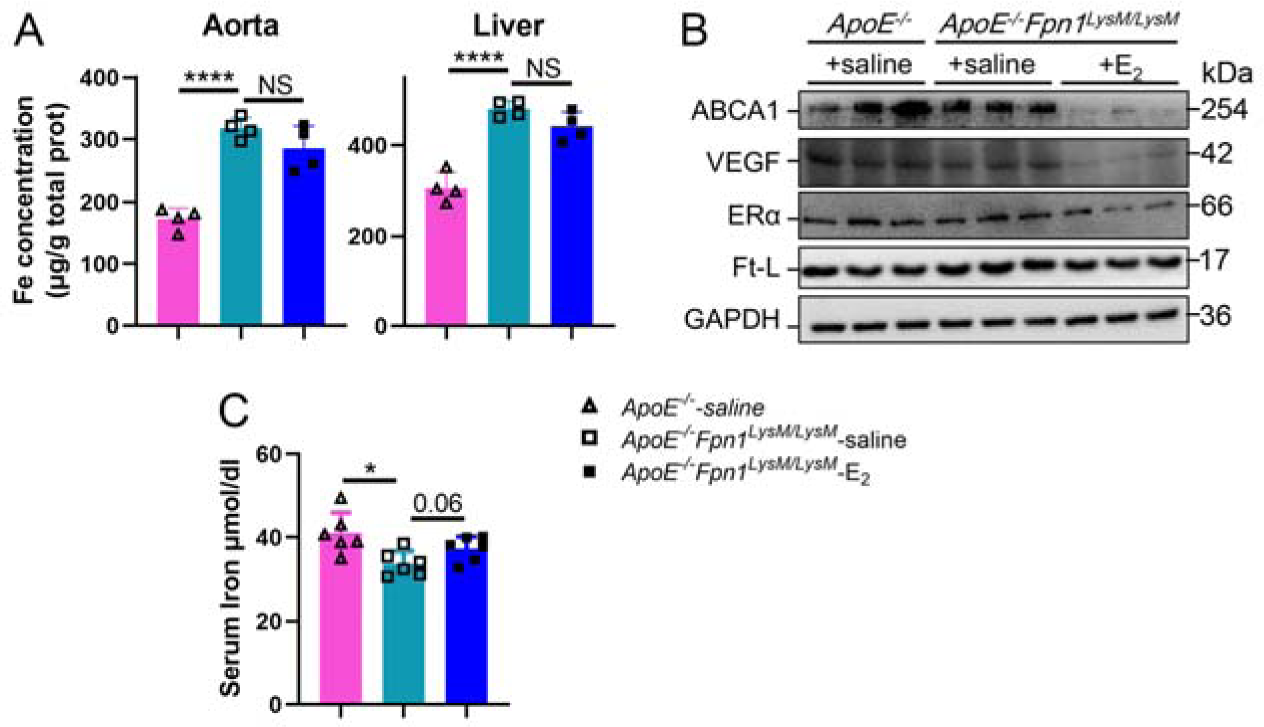
E_2_-triggered ERα deficiency was observed in *ApoE^-/-^ Fpn1^LysM/LysM^* at early post-menopause (25 weeks old). (A) Iron content of aorta and liver detected by ferrozine assays. n = 4, \*\*\*\**p* < 0.0001. (B) ABCA-1, ERα, VEGF and Ft-L protein expression of aorta were detected. (F) Serum iron in different groups. n = 6, \**p* < 0.05.

**Figure S2.**
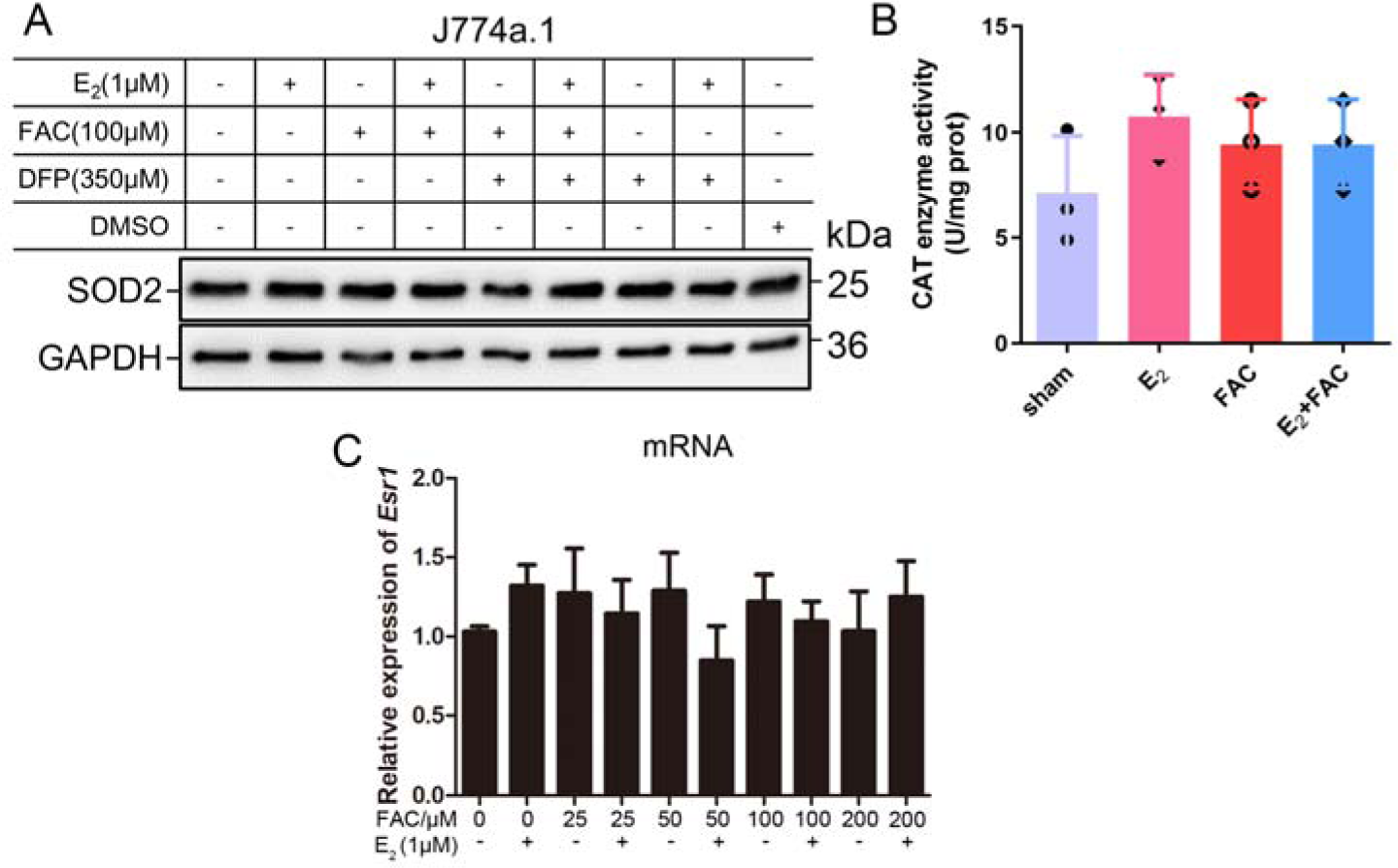
No significant oxidative-stress was raised by application of E_2_ and iron within the indicated concentration. (A) SOD2 protein levels of J774a.1 post treatments with FAC/DFP in the presence/absence of E_2_. (B) The enzymatic activity of catalase in J774a.1. n=4. (C) Relative ERα mRNA expression of J774a.1 treated with different concentrations of FAC in the presence/absence of E_2_, assessed by qPCR. n = 5.

**Figure S3.**
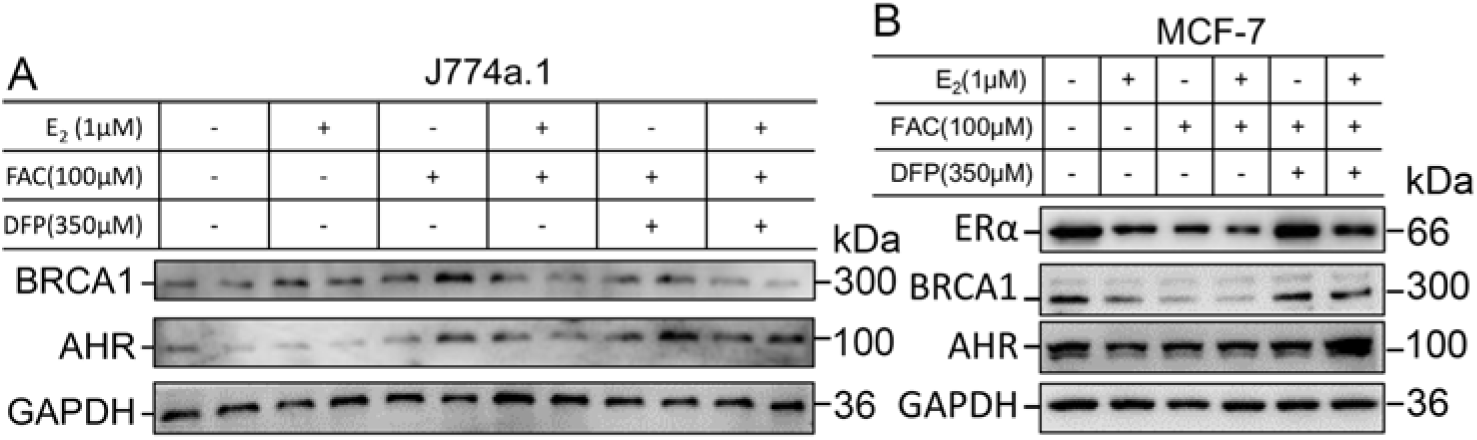
E3-ligase responses to iron and E_2_ treatment in different cell types. (A) BRCA1 and AHR protein expressions in J774a.1 were detected. (B) The protein expression of ERα and its related E3 ligase, BRCA1 and AHR, was detected in MCF-7 cell line. n=4

